# Distinct mechanisms for panoramic and landmark-based view integration in human scene-selective cortex

**DOI:** 10.1101/2025.01.24.634774

**Authors:** Linfeng Tony Han, Russell A. Epstein

**Author notes:** **Correspondence:** Linfeng Tony Han, Russell A. Epstein.

## Abstract

To encode a cognitive map of an environment, a navigating agent must be able to integrate across disparate perceptual views corresponding to the same place. There are two ways that this can be done. First, the agent might integrate across the panorama of views obtainable at a single vantage point. Second, they might integrate across views of a distal location containing a landmark that is visible from multiple vantage points. Guided by previous work, we tested the hypothesis that these two viewpoint-integration processes would be mediated by different neuroanatomical substrates. Male and female human participants were familiarized with a route through a virtual city, along which they closely viewed 24 storefronts that were pairwise associated, either by being located on different buildings directly across the street from each other (same-panorama condition) or by being located on different sides of the same building facing different streets (same-landmark condition). They were then scanned with fMRI while viewing the storefronts in isolation and performing a spatial memory task. Multivoxel pattern analysis revealed a functional distinction between two scene-selective regions: the retrosplenial complex (RSC) showed a significant association effect for same-panorama storefronts, whereas the parahippocampal place area (PPA) showed a significant association effect for same-landmark storefronts. Panoramic association effects were also observed in several other dorsal-stream regions within the medial and lateral parietal lobe. These results demonstrate the existence of two neural mechanisms for integrating across views to represent places as either the observer’s location (same panorama) or the observed location (same landmark).

**SIGNIFICANCE STATEMENT:** An important component of spatial navigation is the ability to integrate across disparate views corresponding to the same place. Here we tested the idea that there are two such integration mechanisms in the human brain. Participants were familiarized with a virtual city and then scanned with fMRI while viewing storefronts from that city. Using multivariate pattern analysis, we find that the retrosplenial complex (RSC) and other parietal regions encode panoramas of views observed from specific locations, whereas the parahippocampal place area (PPA) encodes sets of views corresponding to specific landmarks. These results reveal two anatomically separated mechanisms for integrating views into places, thus advancing our understanding of how the brain forms cognitive maps of spatial environments.

## INTRODUCTION

As we navigate through the world, we experience the environment from multiple points of view, which we piece together into a unified cognitive map (Epstein et al., 2017; O’Keefe & Nadel, 1978; Tolman, 1948). This process requires us to integrate across perceptual representations (“views”) to form abstracted spatial representations (“places”). There are at least two kinds of integration that we can perform. First, integration across the set of views obtained from the same spatial position (i.e., the same vantage point). Second, integration across the set of views corresponding to the same object or location observed from different vantage points. We refer to the first kind of integration as *panoramic integration* and the second kind of integration as *landmark integration*. Both are ecologically important: panoramic integration allows us to recognize our current location and to form a 360-degree mental map of the environment surrounding that location, whereas landmark integration allows us to recognize stable points of reference from a distance and to use these reference points to build up a larger map and orient in space.

Guided by previous work, we hypothesized that panoramic and landmark integration might have different cortical loci within the set of brain regions involved in visual scene processing. Previous fMRI studies have implicated retrosplenial complex (RSC) in panoramic integration by showing neural pattern similarity for images drawn from the same panorama (Berens et al., 2021; Robertson et al., 2016; Vass & Epstein, 2013) and have implicated the parahippocampal place area (PPA) in landmark integration by showing neural pattern similarity for images corresponding to the same landmark (e.g., exteriors and interiors of a building; Marchette et al., 2015). These studies suggest that panoramic and landmark integration have distinct neural bases, centered, respectively, on RSC (panoramic integration) and PPA (landmark integration). However, no previous study has examined both kinds of integration within the same experimental paradigm. The current study was designed to fill this lacuna.

We also set out to address two other issues that remained unresolved from previous work. First, previous studies of viewpoint integration examined learning on different timescales: Robertson et al. (2016) and Berens et al. (2021) examined panoramic associations that were newly learned within an experimental session, whereas Marchette et al. (2015) examined landmark representations constructed over long-term experience. Thus, the previously observed difference in neural locus may be attributable to differences in timescale, rather than differences in the type of integration. Second, previous studies of panoramic integration examined stimuli drawn from real-world environments. While providing ecological validity, such stimuli could not be fully controlled for visual confounds, because adjacent views in real-world environments often share visual features, making it difficult to disentangle whether neural representations are driven by spatial associations or by shared visual characteristics.

To resolve these issues, we developed a 3D virtual navigation paradigm that allowed us to precisely control the temporal and visual properties of the stimuli. Participants were familiarized with a virtual city containing a set of storefronts along a fixed route. The storefronts could be associated in two ways: by being located across the street from each other on different buildings (same-panorama), or by being located on the same building but facing different streets (same-landmark). Participants learned the positions of the storefronts along the route and were then scanned with fMRI while viewing the storefronts in isolation. We obtained storefront-specific neural activation patterns and calculated the similarity between them to test for the coding of panoramic and landmark associations. To anticipate, we observed evidence for panoramic representations in RSC and landmark representations in PPA, consistent with our hypothesis, and we also observed evidence for panoramic representations in other brain regions throughout the dorsal visual processing stream.

## MATERIALS AND METHODS

### Participants

Twenty-four participants (15 female; mean age = 25.9 (± 4.24) years) were recruited from the University of Pennsylvania community. We determined our sample size based on a power analysis based on the effect sizes reported in a similar study on the neural representation of panoramic scene memory (Robertson et al., 2016). These participants were healthy, had normal or corrected-to-normal vision, and were right-handed. Data from five additional participants were excluded before analysis due to MRI facility technical malfunctioning (1 participant), falling asleep in the MRI scanner (1 participant), arriving too late to complete all the MRI scan runs (1 participant), and excessive head motion (2 participants). All participants provided written informed consent in compliance with protocols approved by the University of Pennsylvania Institutional Review Board.

We also recruited 64 participants online from Prolific (prolific.co) for an online testing protocol for selection of stimuli (see *Virtual Environment* section). 7 additional participants entered the online testing protocol, but their data was excluded due to failure in completing the full testing protocol (4 participants) or unusual pattern in response (3 participants).

### Experimental Procedure

#### Virtual Environment

We created a virtual environment in Unity 3D (editor version: 2021.3.2f1, https://unity.com/) that was optimized to test for two kinds of learned spatial associations: panoramic associations and landmark associations. The environment was a virtual city of 400 × 400 virtual meters containing multiple buildings and several streets that crossed at intersections (Fig. 1A). We placed 24 images of storefronts on the sides of the buildings, which could be paired together in two possible ways. Half (12) of the storefronts were paired together by positioning them on different buildings that were on opposite sides of the same street (same-panorama condition, indicated by cyan arrows in Fig. 1A; also see ground-level snapshot in Fig. 1B). The other half (12) of the storefronts were paired together by positioning them on the same building, but on different sides of the building abutting different streets (same-landmark condition, indicated by orange arrows in Fig. 1A; also see ground-level snapshot in Fig. 1B). Four visually salient distal landmarks were located beyond the boundary of the city, one in each cardinal direction (Mountain, Tower, Bridge, and Lighthouse).

**Figure 1.**
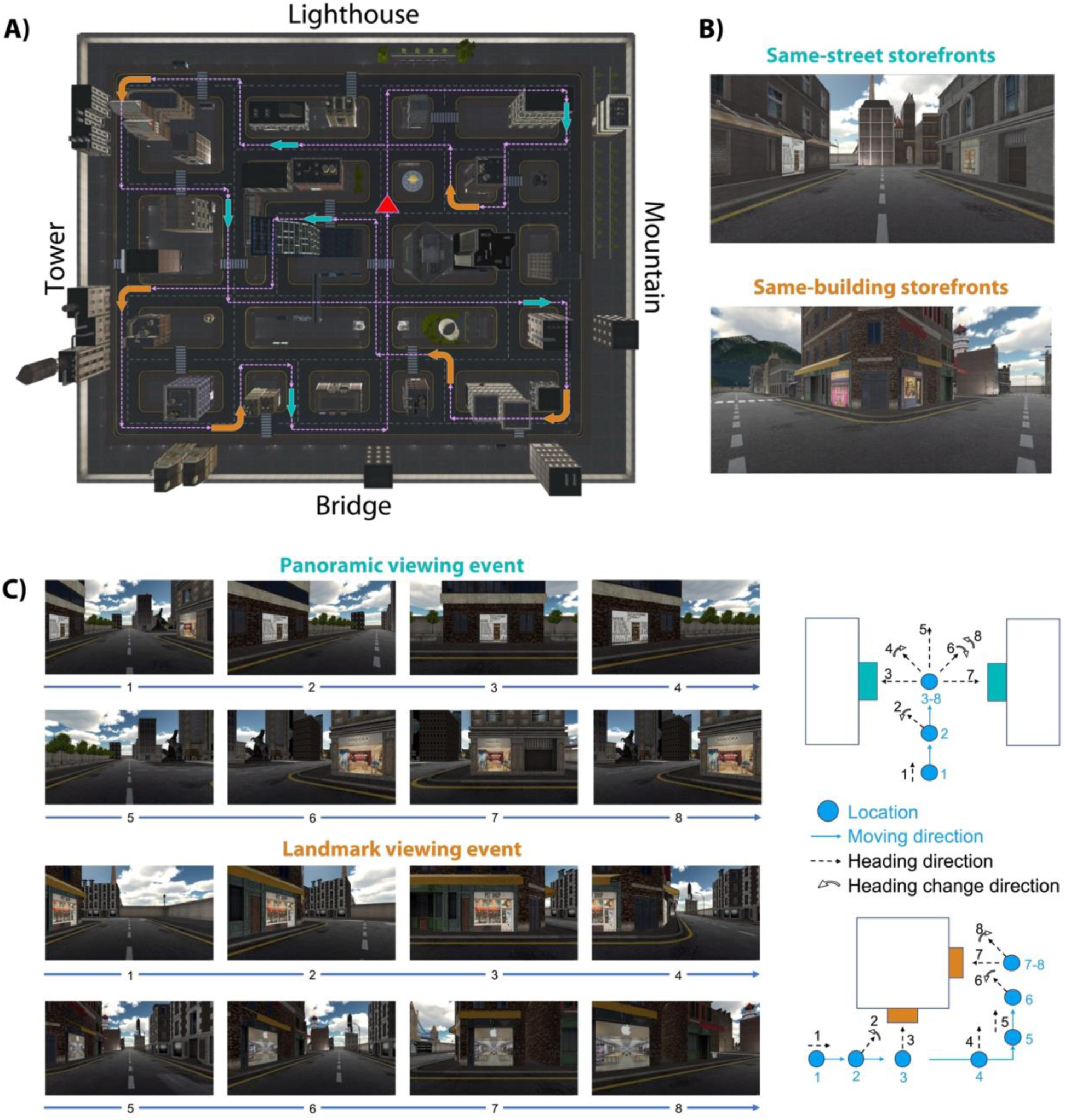
Spatial learning in the virtual city. **A)** Map depicting a bird’s-eye view of the virtual city. Participants viewed a first-person movie showing navigation from ground level along a pre-set route (purple arrows). The route started at the location indicated by the red triangle and returned to the same location at the end. Along the route, participants experienced 12 viewing events. Cyan arrows indicate the route locations of panoramic viewing events during which participants viewed two storefronts located across the street from each other. Orange arrows indicate the route locations of landmark viewing events during which participants viewed two storefronts located on different facades of the same building. Four distal landmarks indicated by words on the map served as directional references. Note that participants never saw the aerial view depicted in the figure. **B)** Ground-level views of the city, showing a same-street storefront pair requiring panoramic integration (top) and a same-building storefront pair requiring landmark integration (bottom). **C)** Sequence of ground-level views (left) and spatial schematic (right) for the panorama and landmark viewing events. Views 1-8 on the left correspond to locations and headings 1-8 on the right. Both events began with the virtual camera progressing straight down the street (1), The camera turned towards the first storefront while continuing to move forward (2), and then paused movement and rotation for 2 s when the first storefront was fully in view (3). In panorama events, the camera then rotated 180 degrees to face the second storefront (4-6), paused for 2 s when it was fully in view (7), and then rotated back 90 degrees to face along the street as forward motion resumed (8). In landmark events, the camera moved sideways from the first storefront until the corner was reached (4), moved forward along the second street (5) while turning towards the second storefront (6), paused for 2 s when the second storefront was fully in view (7), and then rotated back 90 degrees to face along the street as forward motion resumed (8). The two event types were equalized for the time that the storefronts were fully in view (3 and 7) and the temporal interval between these views (4 to 6).

The storefront images were digital photographs of real-world stores that met the following criteria: 1) images were easily classifiable into a store category (e.g., restaurant, jewelry store, bank); 2) images were visually dissimilar from each other. 3) images were easy to memorize and minimally confusable with each other. To meet these criteria, we first collected an initial set of 150 storefront images spanning 22 categories from online sources. We then reduced this set based on responses from raters in an online testing protocol compiled in Qualtrics (https://www.qualtrics.com/) and distributed through Prolific (https://www.prolific.com/). Each rater was presented with half (75) of the storefront images and evaluated the category of each by typing a word or phrase. They were then given a surprise recognition memory test on all 150 storefront images. The images were sequentially presented and rated whether each image had been previously presented on a 4-point scale, choosing from “certainly not,” “maybe not,” “maybe yes,” and “certainly yes.” We also computed the visual similarity across the storefront images based on three commonly used image analysis models: the pixelwise intensity model, the GIST model (Oliva & Torralba, 2001, 2006), and the simple texture model (Renninger & Malik, 2004). Using these behavioral responses and image similarities as a guide, we chose 24 storefront images from 12 categories that maximized categorical distinctiveness and memorability while minimizing visual similarity. The categories were: Bank, bar, barbershop, bookstore, clothing store, electronics store, furniture store, jewelry store, shipping service, pet shop, printing service, and telecommunication service.

We used a virtual environment in this study for two reasons: First, it allowed us to precisely control the temporal interval between the storefronts during learning. Second, it allowed us to control visual similarity among the storefronts. Thus, use of a virtual environment ensured that any neural similarity effects were due to spatial association alone rather than temporal proximity or overlap of visual features.

### Experimental sequence

The experiment was conducted in a single session that includes three separate phases: 1) The spatial learning phase, in which participants navigated through the virtual city and learned the spatial association of the storefronts; 2) The fMRI scanning phase, in which participants performed a judgment of relative direction task on the storefronts, along with several functional localizer tasks; 3) The post-scan behavioral testing phase, in which participants completed a behavioral test that gauged their explicit knowledge of storefront associations.

#### Spatial learning phase

This phase began with participants viewing a short video that familiarized them with the appearance and names of the four distal landmarks surrounding the city. They then completed 8 runs of the spatial learning task, during which they watched a video depicting navigation on a set route through the city. The route passed by all 24 storefronts in a fixed sequence. The route and the storefront locations were identical across all participants, but the assignment of storefront images to locations was randomized across participants to ensure that spatial effects were not attributable to visual, categorical, or other differences between the storefronts. Before entering these spatial learning runs, participants were instructed to try their best to remember (1) the location of each store, (2) the facing direction of each store relative to the distal landmarks, and (3) which stores were across from each other on the same street and which stores were on the same building.

Within each continuous spatial learning run, there were 12 notional viewing events related to the 24 storefronts along the route. In six of the events, participants viewed same-panorama pairs of storefronts located across from each other on the same street, and in the other six events they viewed same-landmark pairs of storefronts located on the same building but facing different streets. For same-panorama events, the virtual camera panned 90 degrees towards the storefront on one side of the street as the point between the two storefronts was approached, paused facing the storefront for 2 s, rotated 180 degrees to face the second landmark, paused facing that storefront, and then rotated 90 degrees back towards the route direction while continuing its progress along the route. For same-landmark events, the virtual camera panned 90 degrees toward the first storefront as the point on the street closest to the storefront was approached, paused facing that storefront, moved around the street corner while maintaining the same facing direction, rotated towards the second storefront as the point on the street adjacent closest to the storefront was approached, paused facing the second storefront, and then rotated 90 degrees back towards the route direction while continuing its progress along the route. The timing was equated across conditions, such that each storefront was directly faced for 2 s, the interval between these direct-facing periods was 5.5 s, and the total duration for all viewing events was 13.5 s. The total continuous on-screen time, including the time that storefronts were coming into view or going out of view, was similar for all storefronts, at 6.0 s (first storefront in panoramic viewing), 5.2 s (second storefront in panoramic viewing), 6.3 s (first storefront in landmark viewing), and 6.3 s (second storefront in landmark viewing), respectively. Detailed procedures of approaching and viewing the two storefronts were depicted in Fig. 1C. Video demos can be found online in our project repository.

To ensure that participants actively engaged in learning the storefronts and their orientations within the environment, a small portion of viewing events were modified to include test questions. In spatial learning runs 2, 3, 6, and 7, these questions probed storefront identify (for 4, 8, 6, and 6 storefronts, respectively, with each storefront tested exactly once), and in spatial learning runs 4, 5, 6, and 7, these questions probed facing direction (for 4, 8, 6, and 6 storefronts, respectively, with each storefront tested exactly once). In identity test events, one of the two storefronts was visually occluded by a gray mask. Upon facing this storefront, participants were presented with small images of three storefronts, and had 15 s to press a button on the keyboard indicating which storefront was behind the occlusion. The visual occlusion disappeared after a choice was made and the viewing event continued as normal. In heading direction test events, a distal landmark’s name appeared on screen after one of the storefronts was approached (the storefront was not occluded). Participants had 15 s to press a button on the keyboard indicating whether the distal landmark was on the front, back, left, or right side relative to their current location. No test questions were included in spatial runs 1 and 8 and participants had an uninterrupted experience of viewing the path through the virtual city.

#### fMRI scanning phase

The objective of the fMRI experiment was to investigate the neural representation of local spatial associations, namely, panoramic and landmark-based associations. To this end, we scanned participants when they performed a variant of the judgment of relative direction (JRD) task (Fig. 2A). On each trial of the JRD task, the participants saw a storefront image in the center of the screen (without any surrounding spatial context) and the name of a distal landmark below the storefront image. The names of the distal landmarks were padded with non-letter characters to eliminate any confounding effects related to the number of letters. Participants were instructed to imagine themselves standing in front of the storefront and directly facing it in the virtual city, and to indicate whether the distal landmark would be to the front, back, left, or right from that imagined viewpoint. Thus, this task required participants to retrieve spatial representations from memory pertaining to the storefront viewed on each trial. However, it did not require them to explicitly recall or report other storefronts that might be associated with the viewed storefront.

**Figure 2.**
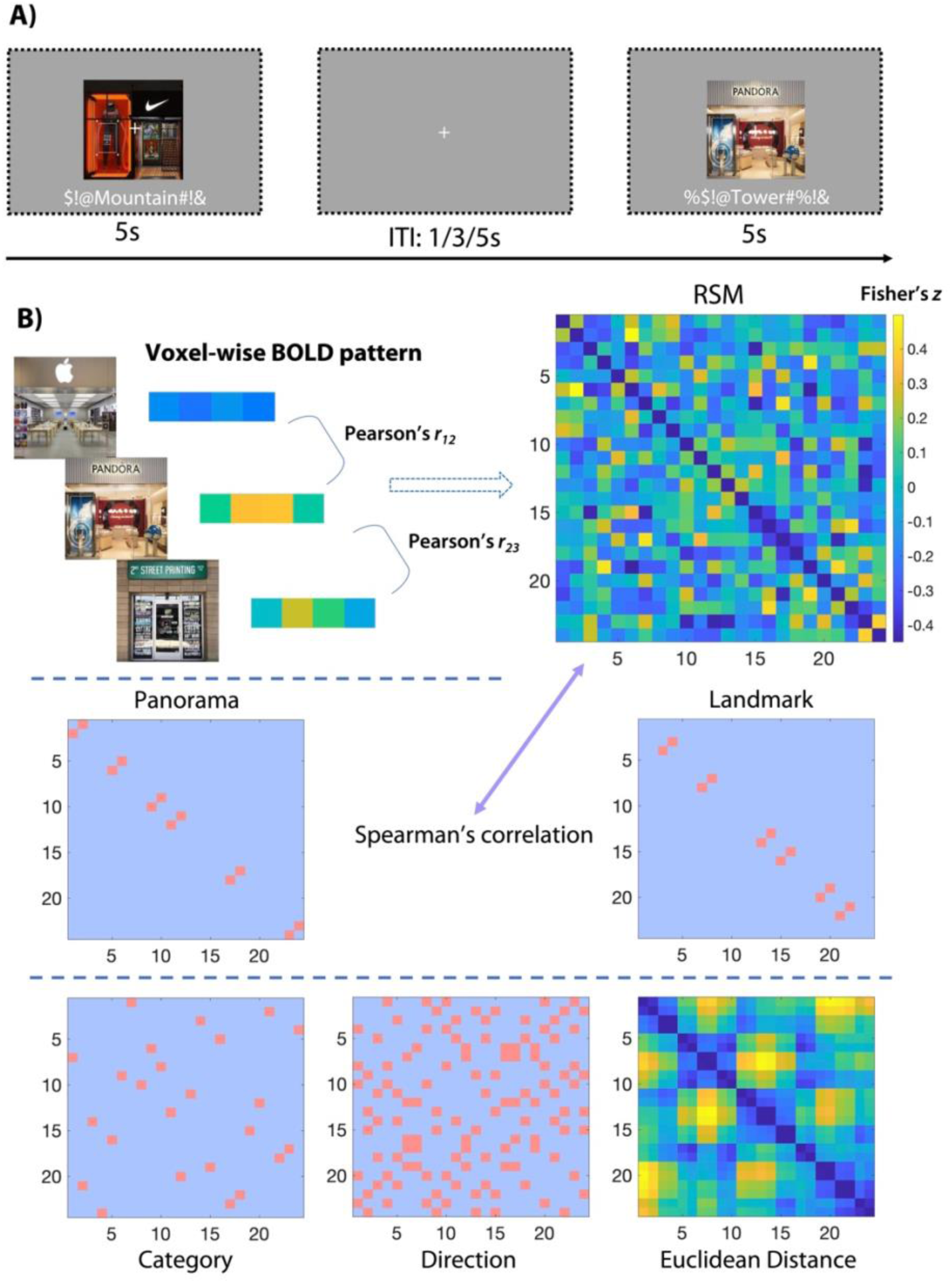
Judgment of relative direction (JRD) task in MRI scanner and representational similarity analysis (RSA). **A)** On each trial of the JRD task, participants saw a storefront image and the name of one of the four distal landmarks. They were instructed to imagine themselves within the city standing in front of the storefront, and to indicate the egocentric direction of the distal landmark from that position and heading. Each trial lasted 5 seconds, with a jittered ITI of 1, 3, or 5 seconds. **B)** To perform representational similarity analyses, a general linear model was used to extract multivariate BOLD activation patterns for the 24 storefronts. Pattern similarities between each pair of storefronts were calculated using Pearson’s correlation and used to construct a representational similarity matrix (RSM) for each region of interest. These RSMs were then correlated with model matrices of interest (panorama, landmark, category, direction, and Euclidean distance) using Spearman’s rank correlation. The RSM on the top-right corner is a sample RSM for one ROI (bilateral RSC) of one participant.

Initial training on the JRD task took place prior to scanning, during the spatial learning phase. Participants completed 24 trials (one trial for each storefront) between spatial learning runs 5 and 6 and another 24 trials between runs 6 and 7. On each training trial, participants had 6 s to answer each question, and no feedback was given. Then, in the fMRI scan session, participants completed 4 runs of the JRD task. Each of the 24 storefronts was presented twice within each scan run, each time in combination with a different distal landmark (48 trials per run). Across pairs of runs (1 and 2, 3 and 4), all the possible 96 storefront-landmark combinations were used. Each JRD trial in the scanner lasted 5 seconds followed by a variable inter-trial interval (ITI) of 1 second (18 trials in each run), 3 seconds (18 trials), or 5 seconds (12 trials). A fixation cross was always visible in the center of the screen, except for the first 12 seconds and last 12 seconds of each run, during which the screen was blank. Total length of each JRD run was 396 seconds (198 TRs).

Participants also completed three functional localizer scan runs. Two of these runs were used to identify voxels that respond selectively during the viewing of visual scenes (scene perception localizer; Epstein & Kanwisher, 1998) and one was used to identify voxels that respond selectively during the recall of familiar places (place memory localizer; Steel et al., 2021). One scene perception localizer run was administered between JRD runs 1 and 2 and the other was administered between JRD runs 3 and 4. The place memory localizer run was administered after JRD run 4.

In the scene perception localizer runs, participants performed a one-back repetition detection task while viewing 16 s long blocks of scenes, faces, objects, and scrambled objects, interspersed with fixation blocks. Each image was shown for 600 ms, followed by a 400 ms interstimulus interval (ISI) and runs were 336 s (168 TRs) in length.

In the place memory localizer, participants were asked to mentally imagine familiar places and people. Prior to the spatial learning phase, participants wrote down 18 names of familiar places and 18 names of familiar people. These names were then presented in the scanner in 30-second blocks, during which 3 place or face names were presented for 7 seconds each followed by a 3-second interstimulus interval. For place names, participants were asked to imagine themselves located in the place and form a mental image of the surrounding environment. For person names, participants were asked to mentally imagine the person’s face. These instructions were identical to Experiment 1 of Steel et al. (2021). The task was administered in one 384 s (192 TR) scan run, which included 12 s of blank screen at the beginning and the end of the scan.

#### Post-scanning behavioral test

Participants were given a storefront association memory test after exiting the scanner to assess their explicit memory for the spatial associations between storefronts. On each trial (18 total), participants were presented with two storefront images simultaneously, which were same-panorama pairs on 6 trials, same-landmark pairs on 6 trials, and non-associated pairs on 6 trials. Each storefront extended approximately 19.5° of visual angle on the screen. Three buttons were also visible on screen: namely “located across from each other on the same street”, “located on the same building”, and “neither.” Participants indicated whether and how the two storefronts were associated by using a mouse click to press one of these three buttons. No feedback was given.

### MRI acquisition and preprocessing

Scanning was conducted at the Hospital of the University of Pennsylvania on a 3T Siemens Prisma scanner equipped with a 64-channel head coil. We acquired T1-weighted images for anatomical localization using 3D magnetization-prepared rapid acquisition gradient-echo pulse sequence (MPRAGE) protocol (repetition time [TR], 2200 ms; echo time [TE], 4.67 ms; flip angle, 8°; voxel size, 1 × 1 × 1 mm; matrix size, 192 × 256 × 160). T2*-weighted functional images sensitive to blood oxygenation level-dependent (BOLD) signals were acquired using a gradient-echo echoplanar pulse sequence (TR, 2000 ms; TE, 25 ms; flip angle, 70°; voxel size, 2 × 2 × 2 mm; matrix size, 96 × 96 × 81). Field mapping was performed after the MPRAGE scan with a dual-echo gradient echo sequence (pulse repetition time [TR], 580 ms; flip angle, 45°; pixel bandwidth, 260; voxel size, 3 × 3 × 3 mm).

MRI data preprocessing was performed using *fMRIprep* 1.2.4 (Esteban et al., 2019). Head motion was corrected within scan run, with all functional images for each participant realigned to the first image and co-registered to the structural image. Functional and T_1_ images were spatially normalized using the following template: ICBM 152 Nonlinear Asymmetrical template version 2009c (Fonov et al. 2009). The B_0_ field maps were applied to correct for distortions in functional images. Functional images from JRD scans were smoothed with a 3mm Gaussian kernel. Functional images from localizer scans were smoothed with a 6mm Gaussian kernel. Image smoothing was performed in SPM12.

### fMRI Data Analysis

#### Estimation of blood-oxygen level dependent responses

For the JRD scan runs, we estimated the voxel-wise blood-oxygen level dependent (BOLD) responses within each scan run to the 24 storefronts using general linear models (GLMs) implemented in SPM12 (Wellcome Trust Centre for Neuroimaging) and MATLAB (R2022b, Mathworks). Trials were modelled as impulse response functions convolved with a canonical hemodynamic response function. Trials with the same storefront image but different distal landmarks were combined into the same condition. Nuisance regressors were used to model the 6 head motion parameters and the global signal in cerebro-spinal fluid (GS-CSF). We then contrasted the beta maps for each storefront against zero to generate t-contrast maps, which were used for further representational similarity analysis.

For the functional localizer scan runs, we followed the same pipeline and estimated the voxel-wise BOLD responses to the stimulus categories (scenes, faces, objects, and scrambled objects for the scene perception localizer; imagined places and imagined faces for the place memory localizer). Each category was modelled as a boxcar regressor convolved with a canonical hemodynamic response function. Nuisance regressors included in the model include included 6 head motion parameters and GS-CSF. The outputs of the GLMs included 2 beta-estimate maps for each stimulus category (one for each run).

#### Regions of interest (ROIs)

Our major goal was to examine the neural representations of panoramas and landmarks in scene-selective regions. To this end, we defined the three standard scene-selective regions—parahippocampal place area (PPA), retrosplenial complex (RSC), and occipital place area (OPA)—in each participant individually using scene-perception functional localizer data and group-level masks derived from a previous study (Julian et al., 2012). We contrasted the beta-estimate maps of scenes against objects, which generated a t-contrast map for scene selectivity for each individual participant. We selected the top 200 voxels with the highest activity within the pre-defined parcel in each hemisphere. The hemisphere-specific ROIs were then combined into bilateral ROIs consisting of 400 voxels. We also defined early visual cortex (EVC) by contrasting the beta-estimate maps of scrambled objects against zero and extracting the top responding 400 voxels in V1 with the constraining mask from Neurosynth (neurosynth.org).

We also defined four place-memory ROIs using data from the place-memory localizer scan. Three of these ROIs were anatomically adjacent to their scene-selective counterparts (PPA, RSC, OPA), consistent with previous descriptions (Steel et al., 2021) and one ROI was overlapping with the superior parietal lobule (see Results for details). Because we had only a single place-memory run for each participant, we defined these regions based on group-level data. For each ROI, we selected 200 voxels in each hemisphere showing greater response to place memory than for face memory. These group masks were applied to all participants in further analyses.

#### Representational similarity analysis (RSA)

We used a standard RSA approach to test for coding of spatial associations and other hypothesized quantities (Fig. 2B). Neural representational similarity matrices (RSMs) were constructed for each ROI in each participant and then compared to model matrices. To construct the neural RSMs, we extracted multi-voxel activation patterns for each of the 24 storefronts based on the t-contrast maps generated across 4 JRD scan runs. The voxel-wise mean activation pattern of all 24 storefronts was then subtracted from each storefront pattern (known as the cocktail mean removal). Pearson’s correlation was calculated between each pair of activation patterns, and these values were Fisher’s z-transformed values to construct the 24 × 24 RSMs. These neural RSMs were then correlated with model RSMs using Spearman’s rank correlation. Each Spearman’s correlation coefficient (Spearman’s *rho*) was transformed to Fisher’s *z* value for averaging across participants.

To test for coding of spatial associations, we used two model matrices that represent panoramic and landmark association, respectively. These were 24 × 24 matrices, with each cell representing the spatial association between each pair of storefronts. For each participant, the model matrix for panoramic or landmark association has 12 cells with 1-values that represent the 6 pairs of storefronts associated either by panorama or landmark. The values for all other cells were zero as the other pairs of storefronts were not locally associated. We correlated the neural RSM for each ROI in each participant with these two model matrices. If a specific ROI contains information about panoramic or landmark association, the correlation between the ROI-specific RSM and the corresponding model matrix should be positive, indicating that same-panorama (or same-landmark) storefronts elicit more similar activation patterns than different-panorama (or different landmark) storefronts.

Next, we extended this step to other spatial and non-spatial quantities: (1) Euclidean distance, modeled as a continuous variable that represents the spatial distance (in virtual meters) between storefronts in the virtual city. (2) Facing direction, modeled as a binary variable that represents whether or not each pair of storefronts shared the same global facing direction. The 24 storefronts were equally grouped into 4 facing directions, so that each storefront shared the same facing direction with 5 other storefronts. (3) Category, modeled as a binary variable that represents whether the two storefronts shared the same visual category. The 24 storefronts were equally grouped into 12 categories, so that each storefront shared the same category with one other storefront.

#### Searchlight analysis

To test for spatial association effects outside the regions of interest, we conducted a whole-brain searchlight analysis. We created a searchlight sphere (radius = 3 voxels) around each voxel in the brain (whole-brain mask: https://neurovault.org/images/396587/). Following the same procedure for the ROI analysis, we correlated the multivoxel activation patterns for each searchlight sphere with the model matrices and assigned the correlational value to the central voxel. The resulting maps of correlational coefficients were smoothed with a 4mm Gaussian kernel. We then tested the significance of the results across participants in the whole brain. The statistics were calculated across participants with Monte-Carlo permutation with threshold-free cluster enhancement (TFCE) and were corrected for multiple comparisons on the whole-brain level. The analyses were implemented in the CosMoMVPA toolbox in MATLAB (Oosterhof, Connolly, & Haxby, 2016).

## RESULTS

### Participants learned the spatial structure of the city and the associations between the storefronts

Prior to scanning, participants viewed movies depicting first-person navigation on a set route through a city that passed by all 24 storefronts. Pairs of these storefronts could be associated together in two ways: by being viewable from the same position on the route (panoramic association), or by being attached to different facades of the same building (landmark association). Behavioral data obtained during spatial learning, during the scan session, and in post-scan testing confirmed that participants successfully learned the spatial structure of the environment and the associations between the storefronts.

In 6 out of the 8 spatial learning runs, the movie sequence was occasionally interrupted by queries about the identity of the upcoming storefront and the facing direction of the camera. Responses to these questions were highly accurate, even in earlier runs where participants had less experience with the virtual city. The average response accuracies for storefront identity questions were 78.1%, 88.5%, 89.6%, and 95.8% in the four spatial learning sessions where these questions were asked (runs 2, 3, 6, and 7); these were all significantly above the 33.3% chance level (*p*s < 0.001; red line in Fig. 3A). The average response accuracies for facing direction questions were 83.3%, 91.7%, 89.6%, and 86.5% in the four spatial learning runs where these questions were asked (4, 5, 6, and 7); these were all significantly above the 25% chance level (*p*s < 0.001; green line in Fig. 3A).

**Figure 3.**
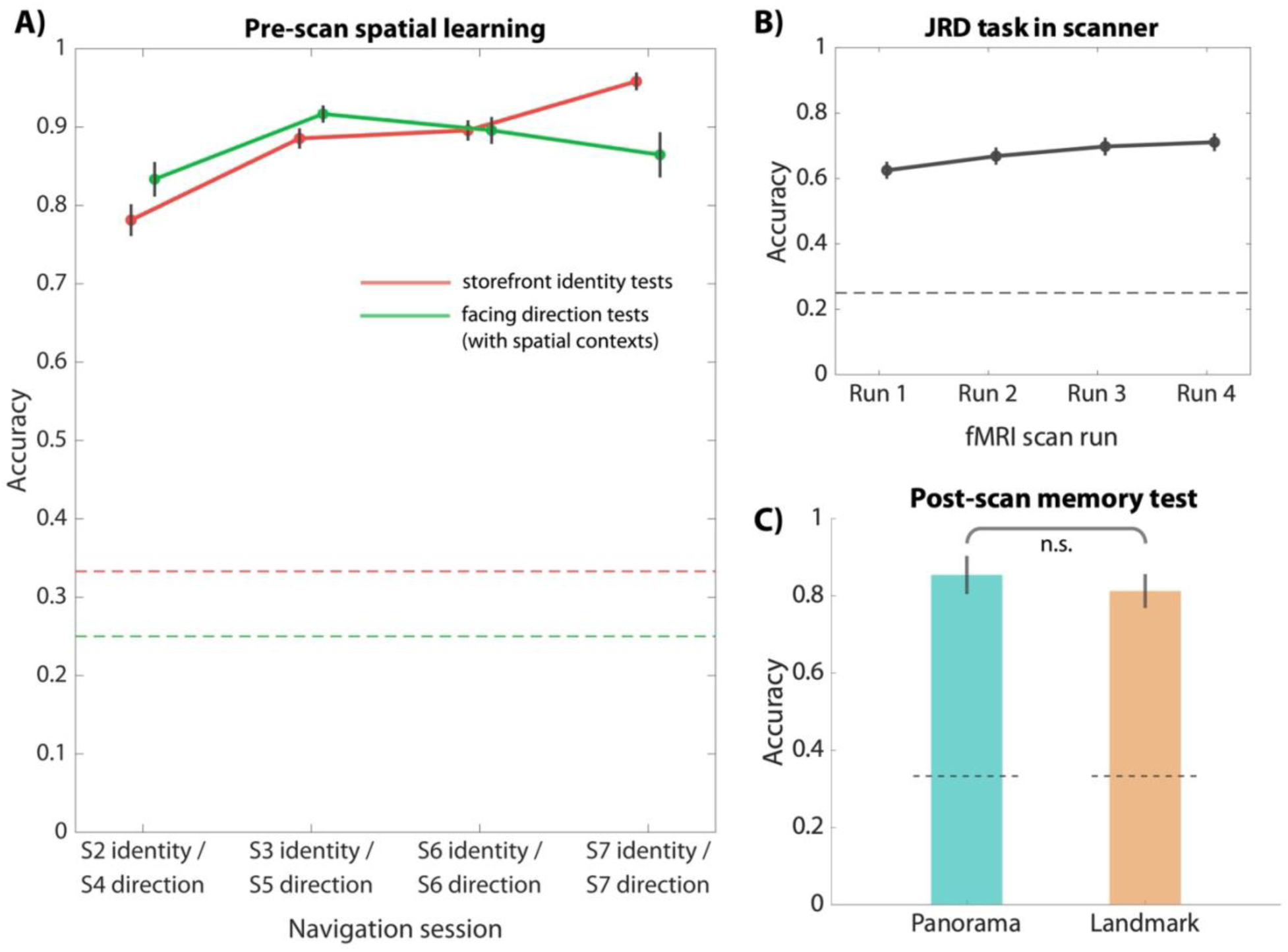
Behavioral performance. **A)** During pre-scan spatial learning, participants were given occasional queries about storefront identity and facing direction. For both types of questions, participants exhibited significantly above-chance behavioral performance, indicating that they learned the location of the storefronts and their headings within the virtual city. Red dashed line indicates the chance level for storefront identity questions (33.3%) and green dashed line indicates the chance level for facing direction questions (25%). **B)** During fMRI scanning, participants completed four scan runs of the JRD task. The group-averaged behavioral performance was significantly above chance (dashed line) for all four scan runs, even though this task was more challenging than the facing direction tests during spatial learning. **C)** After fMRI scanning, participants were administered a pairwise association judgment task, to assess their explicit knowledge of the local spatial association between storefronts. The group-averaged response accuracies for both same-panorama and same-landmark questions were significantly above chance (dashed line). Error bars represent ±1 standard error (SEM).

During scanning, participants performed a JRD task. On each trial, they were shown an image of a storefront and given the name of one of the four distal landmarks. They were asked to imagine that they were standing in front of the storefront and to report whether the distal landmark would be to their left, to their right, in front of them, or behind them from that point of view. No feedback was given. Response accuracies for the four scan runs were 62.5%, 66.8%, 69.8%, and 71.1%, respectively; these were all significantly above the 25% chance level (*p*s < 0.001; Fig. 3B). Performance was lower in this task than in the facing direction questions administered during spatial learning; this was expected as the storefront images were shown outside of the spatial context of the route and a shorter time limit was imposed on each trial. Despite these challenges, participants exhibited good performance that indicated that they had learned the orientations of the storefronts within the spatial framework of the city.

In the post-scan memory test (Fig. 3C), we queried participants on their explicit knowledge of the associations between the storefronts (i.e, whether they were same-landmark or same-panorama pairs). Accuracy for same-panorama questions was 85.4% (±24.2%) and accuracy for same-landmark questions was 81.3% (±21.6%) – both significantly above the 33.3% chance level (*p*< 0.001). There was no significant difference in accuracy between the two types of questions (*t*[23] = 1.297, *p* = 0.207). This demonstrates that participants learned the panoramic and landmark associations between the storefronts to an equivalent degree.

### Multivoxel pattern analyses reveal differential involvement of scene regions in panorama-based vs. landmark-based associations

We then turned to the main question of the study: How does the brain encode associations between stimuli corresponding to the same panorama (i.e., the same viewing location) or the same landmark (i.e., the same viewed location)? To answer this question, we used representational similarity analysis (RSA) to test for similarity of fMRI activation patterns elicited by storefronts corresponding to the same landmark or the same panorama.

We focused our analyses first on PPA and RSC, because these were the regions that most clearly showed these effects in previous studies. We found that multivariate patterns in PPA represented landmark-based associations (*t*[23] = 2.190, *p* = 0.020, one tailed, Cohen’s *d* = 0.447) but not panoramic associations (*t*[23] = −0.448, *p* = 0.671, Cohen’s *d* = −0.092). In contrast, multivariate patterns in RSC represented panorama-based associations (*t*[23] = 2.833, *p* = 0.005, Cohen’s *d* = 0.578) but not landmark-based associations (*t*[23] = 0.812, *p* = 0.213, Cohen’s *d* = 0.166). Although direct contrasts between the effects within each ROI only trended towards significance (PPA-landmark > PPA-panorama effect: *t*[23] = 1.514, *p* = 0.072, Cohen’s *d* = 0.309; RSC-panorama > RSC-landmark effect: *t*[23] = 1.483, *p* = 0.076, Cohen’s *d* = 0.303), a repeated-measures analysis of variance (ANOVA) revealed a significant interaction between ROI (PPA, RSC) and type of association (panorama, landmark), *F*(1, 23) = 6.242, *p* = 0.020, η_p_^2^ = 0.213 (Fig. 4). Thus, when the two ROIs were compared, PPA was found to more strongly represent landmark-based associations, whereas RSC was found to more strongly represent panorama-based associations.

**Figure 4.**
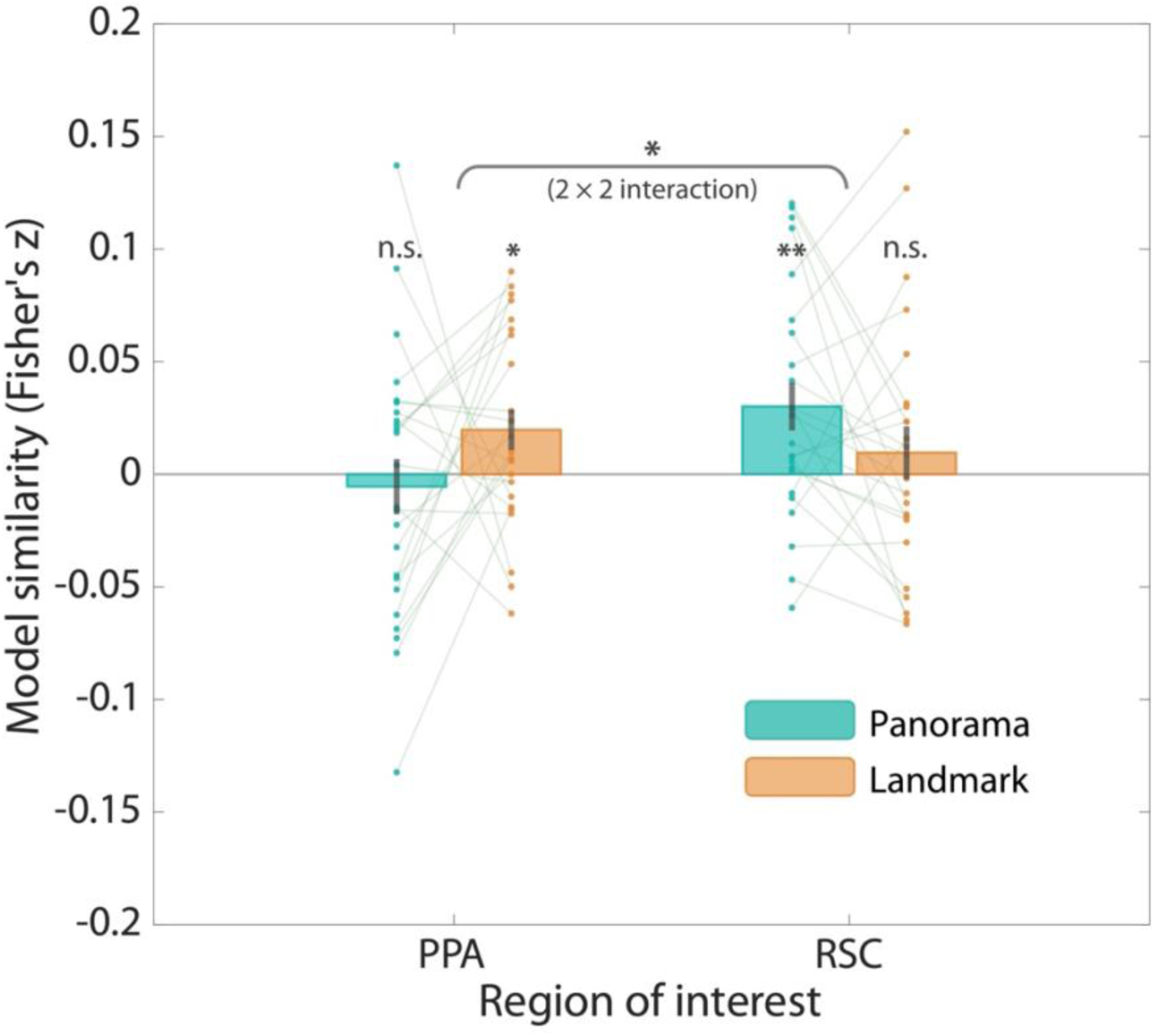
Spatial association representations in PPA and RSC. The RSA fits for each model are plotted. One-sample one-tailed t-tests against zero showed significant coding of panoramic associations in RSC and coding of landmark associations in PPA. A repeated-measures ANOVA revealed a significant 2 x 2 interaction between ROI and type of association. Error bars represent ±1 SEM. Significance markers: \**p*<0.05, \*\**p*<0.01.

Next, we tested the robustness of these effects by examining whether the panoramic and landmark association effects, and the interaction between ROI and type of spatial association, remained significant when across variation in two analysis parameters: the number of voxels included in the ROIs, and whether the mean activation pattern was retained or removed before calculation of pattern similarity (see Materials and Methods, *RSA* section). Mean pattern removal (also known as cocktail mean subtraction) has the potential to systematically adjust multivariate patterns and affect RSA results (Diedrichsen & Kriegeskorte, 2017; Garrido et al., 2013). The selection of ROI sizes may also impact the results or cause biases in interpretation (e.g., Naselaris & Kay, 2015). In our data, the effects of interest remained stable across these analysis parameters (Table 1). We also noticed the following trends: First, retention of the mean pattern made the RSC-panorama effect weaker and the PPA-landmark effect stronger, compared to the results with the mean pattern removed. Second, the RSC-panorama effect increased with ROI size, while the PPA-panorama effect decreased with ROI size.

**Table 1.** Spatial association effects across cocktail-mean removal methods and # voxels selected.

| Cocktail-mean<br>removal | # Voxels in ROI<br>(bilateral) | 2 × 2 Interaction<br>(*F*-statistic) | Panorama (t-statistic) |  | Landmark (t-statistic) |  |
| --- | --- | --- | --- | --- | --- | --- |
|  |  |  | PPA | RSC | PPA | RSC |
| Removed | 100 | 3.438† | 0.050 | 1.749† | 2.772* | 0.808 |
|  | 200 | 3.055† | -0.057 | 1.873† | 2.658* | 0.984 |
|  | 300 | 3.630† | -0.321 | 2.352* | 2.345* | 1.163 |
|  | **400** | **5.853\*** | **-0.448** | **2.833\*** | **2.190\*** | **0.812** |
|  | 600 | 4.105† | -0.204 | 3.495** | 1.234 | 0.669 |
| Unremoved | 100 | 8.135** | -2.118* | 0.960 | 3.800*** | 0.836 |
|  | 200 | 4.857* | -1.955† | 0.966 | 3.377** | 1.248 |
|  | 300 | 4.782* | -2.173* | 1.372 | 3.297** | 1.360 |
|  | 400 | 5.911* | -2.274* | 2.077* | 2.912** | 1.268 |
|  | 600 | 4.304* | -1.614 | 2.413* | 2.069* | 0.937 |

Significance markers: † $p < 0.10$ , \* $p < 0.05$ , \*\* $p < 0.01$ , \*\*\* $p < 0.001$

Beyond PPA and RSC, we also examined association effects within OPA and EVC. OPA is a well-documented scene-selective region that was found to represent panoramic memory in one previous study (Robertson et al., 2016). However, we did not observe any association effects in OPA (panorama effect: *t*[23] = −0.168, *p* = 0.868, Cohen’s *d* = −0.343; landmark effect: *t*[23] = −0.753, *p* = 0.459, Cohen’s *d* = −0.154). We examined EVC as a control region where we did not expect to observe any association effects. Indeed, EVC did not show coding of panorama or landmark (panorama effect: *t*[23] = 0.729, *p* = 0.474, Cohen’s *d* = 0.149; landmark effect: *t*[23] = 0.349, *p* = 0.730, Cohen’s *d* = 0.071).

### Panoramic association effects were observed in dorsal-stream place-memory regions

Beyond the scene-selective regions that were defined by the individual scene-perception localizer data, we also defined place-memory regions using the place-memory localizer data. We examined these regions because previous work has shown that they exhibit systematic network-level functional distinctions with their scene-selective counterparts (Steel et al., 2021), and our experiment taps memory of spatial association that one might expect to be represented in these regions involved in place memory. First, we examined the group data from the place memory localizer to see if we could replicate the same set of “place memory” regions that were identified by Steel et al. (2021). We contrasted place memory against face memory for each participant, and all participants’ contrast maps were entered into a second-level analysis to generate group-level activation maps. This analysis revealed four clusters for place memory (Fig. 5A). These included the ventral place-memory area (VPMA), medial place-memory area (MPMA), lateral place-memory area (LPMA), as described in Steel et al. (2021). We also observed a fourth place memory region within the superior parietal lobule. This region was observed in the previous study (Steel et al., 2021, Fig. 2), but it was not commented upon or named; here we label it the superior parietal place-memory area (SPPMA). The first three clusters (VPMA, MPMA, and LPMA) were adjacent to their scene-perception counterparts (PPA, RSC, and OPA), with a high degree of overlap in the case of PPA/VPMA and RSC/MPMA.

**Figure 5.**
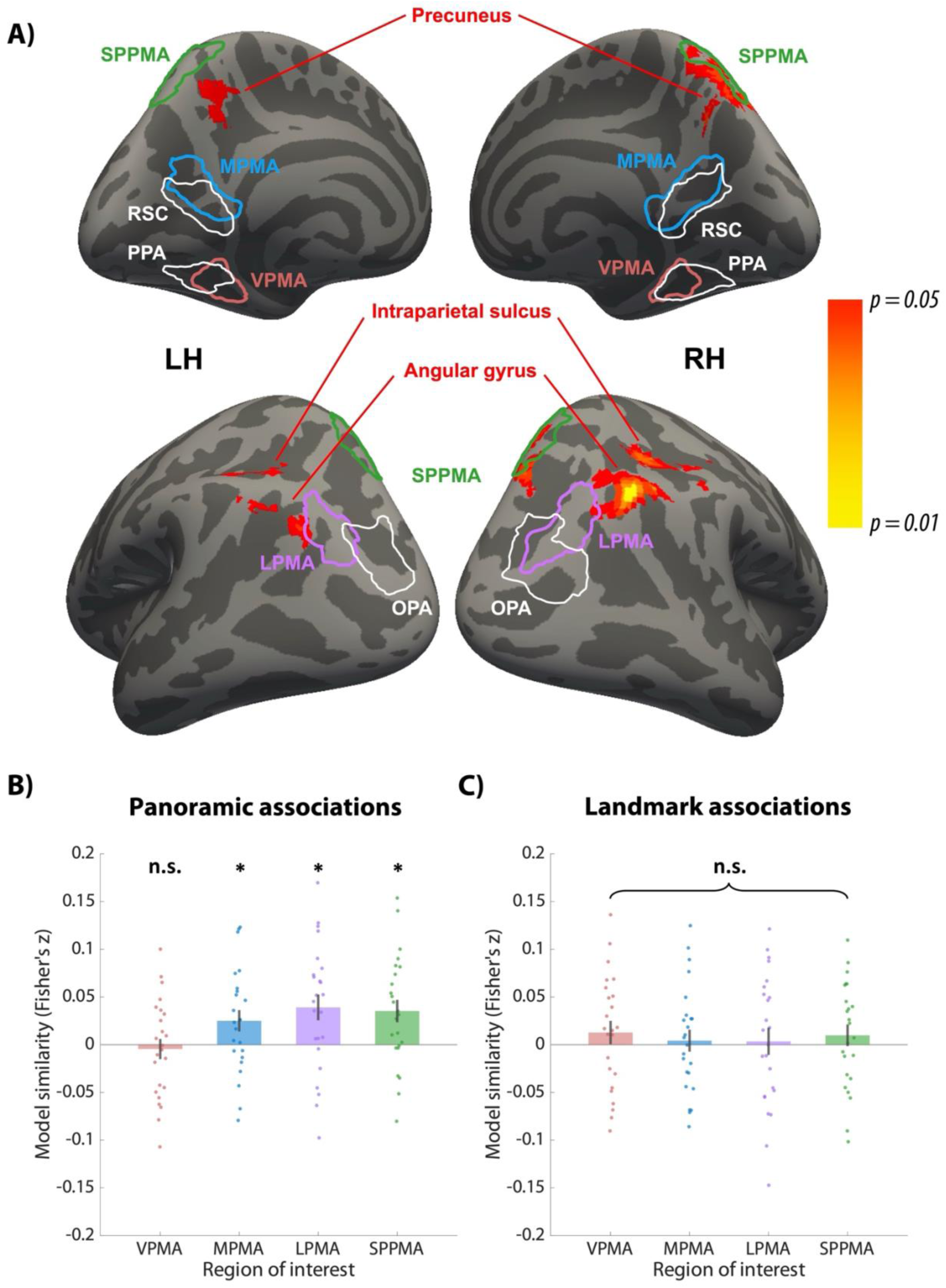
Place-memory ROIs and whole-brain searchlight results. **A)** Anatomical locations of place-memory and scene-selective regions, overlaid with searchlight activation map for panoramic associations. Place-memory regions were defined based on place-memory localizer scan. For visualization purposes, group-wise place memory > face memory results are thresholded at uncorrected *p* < 10^-5^, exhibiting four prominent clusters: VPMA, MPMA, LPMA, and SPPMA (outlined in red, blue, purple, and green, respectively). VPMA, MPMA, LPMA were anatomically adjacent to their scene-selective counterparts: PPA, RSC, and OPA. 200 voxels with the highest group-wise activity level in each hemispheric cluster were selected and combined bilaterally for ROI analysis. Several parietal lobe regions showed significant coding for same-panorama views in the searchlight analysis, including the angular gyrus, superior parietal lobule, intraparietal sulcus, and precuneus. The searchlight map was thresholded at *p* < 0.05 after permutation testing and correction for multiple comparisons across the whole brain. **B)** MPMA, LPMA, and SPPMA showed significant coding of panoramic associations. Results are corrected for multiple comparison across 4 ROIs. Significance marker: *0.01<*p_corrected_*<0.05. **C)** No place-memory regions showed significant coding of landmark associations (all *p*s > 0.150).

For each of the four place memory regions, we identified the 200 top responding voxels in each hemisphere on the group level and combined across hemispheres to construct a 400-voxel ROI. We then tested whether the activation patterns in these ROIs contained information about panoramic and landmark associations. We had no *a priori* hypotheses about the effects that we expected to find, so for each kind of spatial association we applied Holm-Bonferroni correction for across 4 ROIs to avoid false positives. MPMA, similar to RSC, showed significant coding of panoramic associations (*t*[23] = 2.232, *p_corrected_* = 0.036, Cohen’s *d* = 0.456, Fig. 5B, blue bar), which is not surprising given the substantial anatomical overlap between MPMA and RSC. LPMA also showed significant coding of panoramic associations (*t*[23] = 2.896, *p_corrected_* = 0.012, Cohen’s *d* = 0.591, Fig. 5B, purple bar), which contrasts to the lack of a panoramic coding effect in the adjacent but largely non-overlapping OPA. SPPMA also showed significant coding of panoramic associations (*t*[23] = 2.992, *p_corrected_* = 0.013, Cohen’s *d* = 0.611, Fig. 5B, green bar). None of these place-memory regions exhibited significant coding of landmark associations, including VPMA, which overlaps with PPA (Fig. 5C, all *p*s > 0.150). The difference between the PPA and VPMA results might be because PPA was defined individually for each participant whereas VPMA was defined at the group level.

### Searchlight analyses revealed panoramic association effects in several parietal lobe regions

We also performed a searchlight analysis to test for spatial association effects outside of the predefined ROIs. The results were corrected for multiple comparisons across the whole brain, and thus required a stronger underlying effect than in the ROI analyses to reach significance. No regions showed landmark-based association effect at this threshold. However, the panoramic association effect was observed in several parietal regions (Fig. 5A). One of these regions was in the superior parietal lobe and overlapped substantially with SPPMA, one of the place-memory areas identified with our place-memory localizer. Significant panoramic association effects were also found in the angular gyrus, intraparietal sulcus, and precuneus.

### Coding of other spatial and non-spatial quantities

Finally, we examined whether the scene-selective and place-memory regions of interest represent other spatial and non-spatial quantities including Euclidean distance, facing direction, and storefront category (Fig. 6). We submitted the RSA for four place-memory ROIs, three scene-selective ROIs, and EVC. We did not have *a priori* hypotheses for these effects, so to avoid false positives, we applied Holm-Bonferroni correction across all 8 ROIs. Results showed significant coding of Euclidean distance between storefronts in LPMA (*t*[23] = 4.327, *p_corrected_*= 0.002, Cohen’s *d* = 0.883) but no other regions. Coding of facing direction (which was defined by the distal landmarks in the virtual city) was not significant in any of these ROIs after Holm-Bonferroni correction; only VPMA showed a trend towards significance (*t*[23] = 2.313, *p_raw_* = 0.015, *p_corrected_* = 0.120, Cohen’s *d* = 0.472). Coding of storefront category was observed in several regions: RSC (*t*[23] = 2.743, *p_corrected_* = 0.035, Cohen’s *d* = 0.621), OPA (*t*[23] = 4.445, *p_corrected_* < 0.001, Cohen’s *d* = 0.997), EVC (*t*[23] = 4.135, *p_corrected_* = 0.001, Cohen’s *d* = 0.990), and MPMA that showed a trend towards significance after Holm-Bonferroni correction for 8 comparisons (*t*[23] = 1.899, *p_raw_* = 0.035, *p_corrected_* = 0.176, Cohen’s *d* = 0.424). Notably, category coding was not observed in PPA, in contrast to previous results (Persichetti & Dilks, 2019).

**Figure 6.**
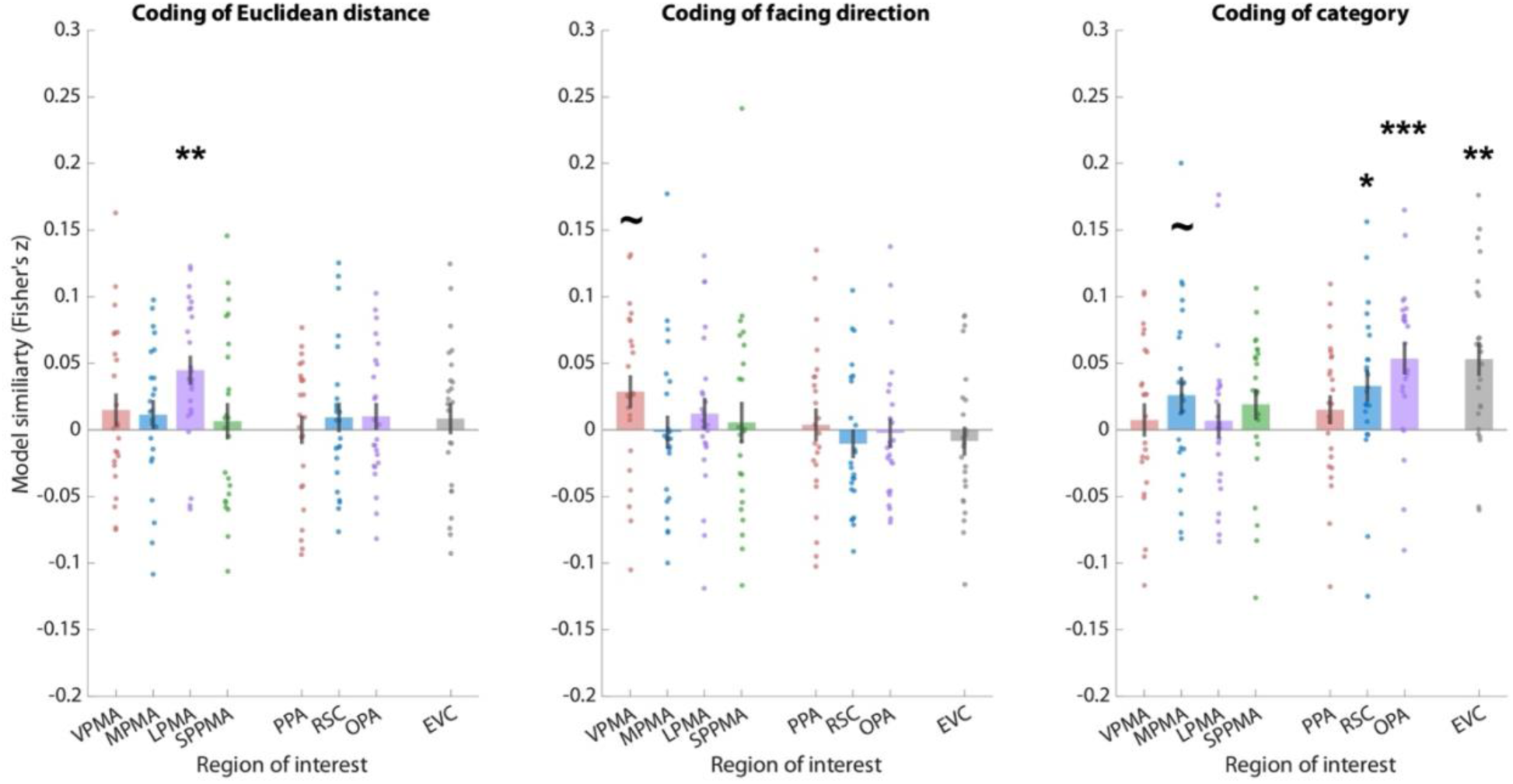
Coding of Euclidean distance, facing direction, and storefront category in scene-selective and place-memory ROIs. Euclidean distance between storefronts in the virtual city is represented in LPMA. There is a trend towards coding of allocentric facing direction defined by distal landmarks in the virtual city in VPMA. Storefront category is represented in MPMA, RSC, OPA, and EVC. Results are Holm-Bonferroni corrected for multiple comparison across 8 ROIs. Significance markers: \**p*<0.05, \*\**p*<0.01, \*\*\**p*<0.001, ∼*p_raw_*<0.05, but insignificant after Holm-Bonferroni correction.

## DISCUSSION

The goal of this study was to understand how the human brain integrates across multiple views corresponding to the same “place.” We examined two kinds of integration: integration across views *acquired from* the same location (panoramic integration) and integration across views *directed at* the same location (landmark integration). We tested these two kinds of integration by familiarizing participants with storefronts located along a route within a virtual city, and then scanning them with fMRI while they viewed the storefronts in isolation and performed a JRD task. Within the city, the storefronts could either be located across the street from each other (same panorama) or on different sides of the same building (same landmark). We found that multivoxel activation patterns in RSC were more similar for same-panorama storefronts (compared to the baseline similarity between unrelated storefronts), whereas multivoxel activation patterns in PPA were more similar for same-landmark storefronts. These results demonstrate that RSC is more strongly involved in panoramic integration while PPA is more strongly involved in landmark integration.

These findings align with prior results implicating RSC in panoramic integration (Berens et al., 2021; Robertson et al., 2016; Vass & Epstein, 2013) and PPA in landmark integration (Marchette et al., 2015; Vass & Epstein, 2017) but go beyond them in several ways. First, previous studies examined either panoramic integration or landmark integration separately, whereas here we examine both kinds of integration within the same experimental paradigm, allowing them to be directly compared. Second, previous studies focused on either spatial representations that were newly learned during the experimental session (e.g., Berens et al., 2021; Robertson et al., 2016) or spatial representations that were formed over long-term, real-world experience (e.g., Marchette et al., 2015). This left open the possibility that previously observed distinctions between RSC and PPA were due to differences in temporal scale, rather than to differences in type of integration. Here we can reject this alternative hypothesis because panoramic and landmark integration were examined together with temporal factors strictly controlled. Third, previous studies of panoramic association used learning paradigms in which panoramas were learned individually, outside of a larger navigational context. Here we used a route learning paradigm within an extended environment that more closely resembles the dynamic, continuous, and immersive nature of real-world spatial experience. Finally, we chose our visual stimuli to minimize confounding effects of visual similarity. This allowed us to disentangle abstract spatial representations from low-level perceptual confounds, an issue that has complicated the interpretation of findings in prior studies using visual snapshots drawn from real-world neighborhoods.

The PPA findings add to the body of literature linking this region to the encoding of landmarks. Early fMRI studies showed that PPA responds preferentially to visual scenes (Epstein & Kanwisher, 1998) and buildings (Aguirre et al., 1998). Subsequent work demonstrated that PPA responds to non-scene and non-building objects based on the navigational significance of the object (Troiani et al., 2014), with stronger activity to objects at decision points compared to non-decision points (Janzen & van Turennout, 2004; Schinazi & Epstein, 2010), larger objects compared to smaller objects (Konkle & Oliva, 2012; Julian et al., 2017), spatially stable objects vs. spatially unstable objects (Mullally & Maguire, 2011), and distant objects compared to proximal objects (Amit et al., 2012). Using multivoxel pattern analysis, Marchette et al. (2015) showed that PPA represents associations between the interior and exterior of the same building, whereas Sun et al. (2021) showed that PPA represents associations between landmark objects located in the same interior room. In contrast, Persichetti and Dilks (2019) found that PPA did *not* encode associations between *different* building*s* that were merely proximal in space. Together with the current results, this literature implicates PPA in the coding of individual landmarks, by which we mean distinct items that are uniquely associated with specific spatial locations (including scenes, which are images of the locations themselves). Notably, whereas many of the aforementioned studies found landmark effects in several scene regions, here we find landmark effects in PPA alone. This may be because the current paradigm strictly controlled for visual and physical differences between the stimuli, allowing the landmark association effect to be more purely isolated. The RSC results extend previous work implicating this region in the encoding of panoramas. Park and Chun (2009) observed adaptation in RSC (but not PPA) when participants were shown sequences of images corresponding to adjacent views from a visual panorama. Vass and Epstein (2013) found that multivoxel patterns in RSC were more similar for panoramic views taken at the same familiar campus intersection than for views taken at different campus intersections, whereas Robertson et al. (2016) observed that multivoxel patterns in RSC (and OPA) were more similar for endpoint views of the same panorama if participants had previously experienced the portion of the panorama connecting the two views. Berens et al. (2021) observed a similar panorama effect in both RSC and PPA but found that only the RSC effect was predictive of participant’s ability to identify the two views as the same location. Our study replicates and extends these findings by introducing more controlled experimental conditions, demonstrating that RSC robustly encodes panoramic associations in dynamic spatial navigation and even when visual confounds are minimized. These results reinforce the notion that RSC supports the integration of fragmented visual inputs into unified spatial representations centered on the observer.

Beyond PPA and RSC, we found additional regions in the dorsal stream involved in panoramic integration. ROI analysis revealed significant coding of panoramic associations in three place memory areas (MPMA, SPPMA, LPMA). Whole-brain searchlight analyses revealed panoramic association effects in several additional parietal lobe regions, including the angular gyrus, precuneus, and intraparietal sulcus. These parietal regions are known to encode self-related spatial information (Boccia et al., 2014; Ciaramelli et al., 2010; Murias et al., 2019) and have extensive connectivity with RSC (see Alexander et al., 2023 & Chrastil, 2018; for reviews). These results suggest the encoding of representation tied to the observer’s location involves a network of regions in the medial and lateral parietal lobe, including RSC but extending beyond it.

Our investigation into other spatial and non-spatial quantities yielded additional insights into the functional roles of scene-selective and place-memory regions. Coding of Euclidean distances between storefronts was observed in LPMA, suggesting that this region may contribute to representing continuous spatial metrics in addition to representing discrete associations. Coding of store category was observed in RSC/MPMA, OPA, and EVC, but not in PPA. This anatomical distribution is partially consistent and partially inconsistent with previous findings (c.f., Walther et al., 2009; Epstein & Morgan, 2012; Persichetti & Dilks, 2019). We suggest that the anatomical locus of “category” information is likely to vary across studies depending on the visual features that distinguish between categories in each particular case.

Our findings conceptually align with the well-established functional division between the dorsal and ventral visual streams. We found stronger dorsal representations for panoramic associations and stronger ventral representations for landmark associations. The dorsal stream, specialized for “where” processing, is well-suited for encoding panoramic information tied to the observer’s perspective (Goodale & Milner, 1992; Mishkin & Ungerleider, 1982). Conversely, the ventral stream, which processes “what” information, underpins landmark-based representations by linking spatially significant objects to their context (Bar & Aminoff, 2003; Janzen & van Turennout, 2004; Peelen et al., 2024). This conceptual distinction offers a framework for understanding how the brain integrates diverse spatial cues into cohesive mental maps, with implications for a wide range of real-world navigational tasks.

In conclusion, we demonstrate evidence for two distinct brain mechanisms for integrating separated views into coherent space. Panoramic integration allows the formation of place representations related to the location of the observer, whereas landmark integration allows the formation of place representations related to the location of a stable external entity such as an object or building. Both kinds of place representations are important in day-to-day navigation, where it is important to be able to recognize one’s current location and also to recognize stable points of reference from a distance.

## Acknowledgements

This work was supported by NIH grant R01EY022350. We thank Tamar Japaridze and Marlie Tandoc for assistance with fMRI data collection.

